# GSAP modulates *γ*-secretase specificity by inducing conformational change in PS1

**DOI:** 10.1101/461061

**Authors:** Eitan Wong, George P. Liao, Jerry Chang, Peng Xu, Yue-Ming Li, Paul Greengard

## Abstract

The mechanism by which GSAP (γ-secretase activating protein) regulates γ-secretase activity has not yet been elucidated. Here, we show that knockout of GSAP in cultured cells directly reduces γ-secretase activity for Aβ production, but not for Notch1 cleavage, suggesting that GSAP may induce a conformational change contributing to the specificity of γ-secretase. Furthermore, using an active site directed photoprobe with double cross-linking moieties, we demonstrate that GSAP modifies the orientation and/or distance of PS1-NTF and PS1-CTF, a region containing the active site of γ-secretase. This work offers insight into how GSAP regulates γ-secretase specificity.

## Introduction

γ-secretase is a large intramembrane protein complex comprised of four essential components, Presenilin 1 (PS1), Nicastrin (NCT), Anterior pharynx-defective 1 (Aph1) and Presenilin enhancer 2 (Pen2) ^1,2^. PS1 is the catalytic core of the complex ^3-5^, and requires an activation step by endoproteolysis ^6^ that is dependent on Pen-2.^5,7,8^. In addition to these mandatory subunits, γ-secretase is also regulated by nonessential proteins, such as CD147, TPM21, the γ-secretase activating protein (GSAP) and Hif-1α ^9-12^. GSAP is a ~98 kDa holoprotein, which undergoes extensive processing resulting in a ~16 kDa C-terminal fragment ^12^. This fragment was found to form a ternary complex with γ-secretase and βCTF and to regulate the cleavage at the γ-sites, but not ∈-sites, controlling the generation of Aβ and AICD, respectively. Decreasing GSAP expression in cells significantly reduced Aβ levels ^12,13^ without affecting the cleavage of other substrates, such as Notch ^12^. However, the role of GSAP in the regulation of γ-secretase has been challenged ^13^ due to lack of direct mechanistic evidence. Nevertheless, GSAP RNAi mice crossed with double transgenic APPsweXPS1ΔE9 AD model mice have reduced Aβ burden ^12^, indicating that GSAP is a novel therapeutic target for the treatment of AD. Furthermore, genetic studies have shown that GSAP is linked with aging, AD and Down syndrome ^14-16^. Therefore, investigating the function of GSAP in Aβ production and γ-secretase activity and specificity is critical for developing an in-depth understanding of γ-secretase modulation and effective AD therapeutics. However, such studies have been hindered due to lack of suitable technologies.

## Results and Discussions

Previous studies examined the function of GSAP using siRNA knockdown methods ^12,13^. To further investigate the role of GSAP in γ-secretase regulation, we knocked out (KO) GSAP with CRISPR/Cas9 technology in HEK293 cells that stably express APP (HEK-APP). KO of GSAP was confirmed by genomic sequencing (data not shown). mRNA levels of GSAP in GSAP-KO cells were not detectable (Fig. 1A), whereas the expression of APP mRNA was not changed (data not shown). Secretion of Aβ40 and Aβ42 in the GSAP-KO cells was only 75% and 73% of HEK-APP WT cells, respectively (Fig. 1B and C).

**Figure 1.**
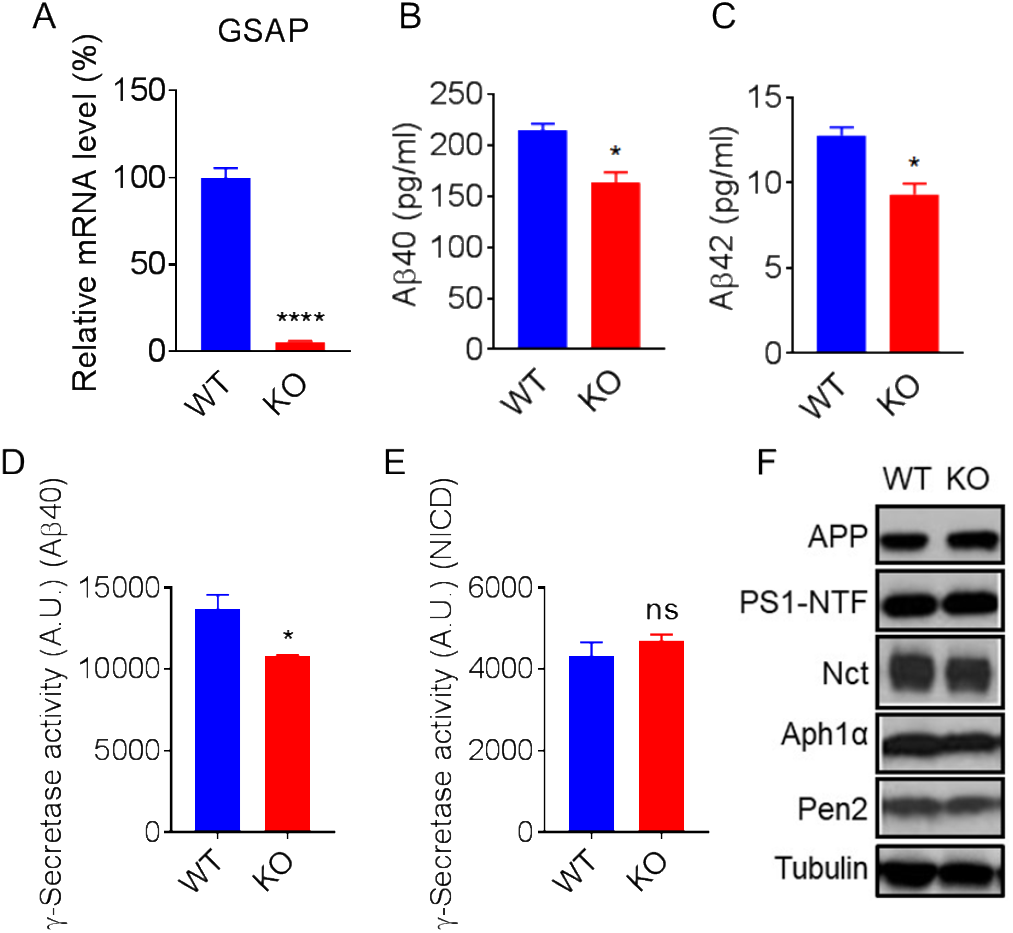
HEK-APP GSAP KO reduces Aβ secretion and γ-secretase activity for APP without changing γ-secretase components. GSAP mRNA levels in HEKGSAP WT versus GSAP KO cells (A). Secretion of Aβ40 (B) and Aβ42 (C) in HEK-APP GSAP WT versus HEK-APP GSAP KO cells showing significant reduction in KO cells. All data represent means ±SEM; n=3. (D) γ-Secretase activity levels toward recombinant APP are decreased in GSAP KO HEKAPP compared to WT counterpart. (E) γ-Secretase activity levels toward recombinant Notch remain the same in GSAP KO HEKAPP compared to WT counterpart. All data represent means ±SEM; n=6. (F) Western blot analysis of APP and total γ-secretase subunits PS1-NTF, Nct, Aph1a, and Pen2 in HEK-APP WT and KO. (*p-value < 0.05, **** < 0.0001)

To directly measure the effect of GSAP on γ-secretase activity, we performed exo-cell assays ^17^ using recombinant APP or Notch substrate ^18^, which allows for the immediate and real-time analysis of γ-secretase activity for both substrates. HEK-APP WT and GSAP KO cells were seeded in a 96-well plate overnight. The recombinant substrates were then added to the cells in the presence of CHAPSO and incubated for 2.5 hours to measure γ-secretase cleavage. The cleaved products were detected with an AlphaLISA assay ^18^. γ-Secretase activity was calculated by normalizing to protein concentration. HEK-APP GSAP KO cells have only 71 % γ-secretase activity for Aβ40 production compared to WT (Fig. 1C), but the same level of Notch1 cleavage (Fig. 1D), indicating that GSAP solely impacts the processing of APP without affecting Notch processing. Next, we tested whether the reduced γ-secretase activity and Aβ secretion were a result of changes in expression levels of the γ-secretase subunits. The protein levels of APP, PS1-NTF, Nct, Aph1a and Pen2 remained unchanged in HEK-APP KO compared to WT (Fig. 1E), indicating that GSAP-KO directly affects γ-secretase activity without altering overall steady-state levels of γ-secretase subunits. The same conclusion was reached when we assayed cell membranes prepared from both WT and KO cells (data not shown). In addition, we generated GSAP-KO SH-5YSY cells and found that GSAP-KO reduces γ-secretase activity for Aβ40, but not for Notch cleavage (Fig. S1).

To determine whether the reduction of γ-secretase activity in the GSAP KO cells directly results from elimination of GSAP, we performed rescue studies by overexpressing GSAP in the KO cells. Empty vector (EV) or the full-length human GSAP with a C-terminal HA tag (hGSAP) construct was transfected into the HEK-APP WT or GSAP KO cells and secreted Aβ species were measured 48 hours post transfection. We found that overexpression of hGSAP in HEK-APP KO cells can fully, or partially restore secreted Aβ40 and Aβ42 (Fig. 2A and B). However, overexpression of hGSAP in WT HEK-APP cells had no effect on Aβ secretion, indicating that the endogenous level of GSAP is sufficient for γ-secretase activation for the processing of APP. Next, we measured γ-secretase activity using cell membranes prepared 48 hours post transfection. In agreement with the Aβ production data, expression of hGSAP in KO cells rescued γ-secretase activity for APP, but the γ-secretase activity remained unchanged in the WT membrane (Fig 2C). Remarkably, γ-secretase activity for cleavage of recombinant Notch substrate was not changed with GSAP overexpression in any of the cell lines (Fig. 2D). Expression of hGSAP, migrating as expected at ~98 kDa protein, was confirmed by Western blot using anti-HA antibodies (Fig. 2E). Moreover, the overexpression of GSAP did not alter the expression of APP or PS1 (Fig. 2E). Similar results were obtained rescuing the γ-secretase activity and Aβ secretion with SH-5YSY GSAP-KO cells (Fig. S2). The expression of GSAP on the GSAP-KO background can rescue γ-secretase activity for the processing of APP, demonstrating that the GSAP-KO effect on γ-secretase activity comes from the abolition of GSAP.

**Figure 2.**
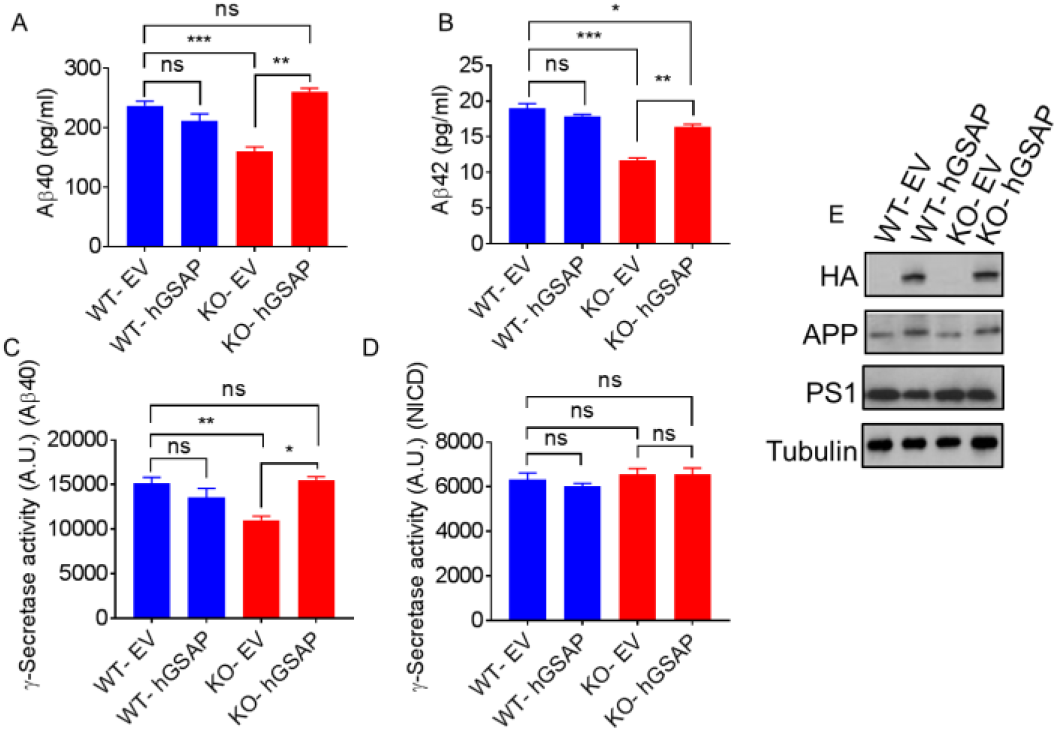
Overexpression of hGSAP in GSAP KO cells rescues γ-secretase activity and Aβ secretion. Empty vector (EV) or full-length human GSAP (hGSAP) were transfected in the HEKAPP GSAP WT or KO cells. Aβ secretion was measured 48 hours post-transfection using MSD Aβ detection; Aβ40 (A) and Aβ42 (B). Membrane fractions prepared from 48 hours post transfection of EV and hGSAP in HEKAPP WT or KO were assayed for γ-secretase activity using recombinant APP (C) or Notch (D). All data represent means ±SEM; n=3 (E) Western blot analysis of HA (hGSAP), APP and PS1-NTF in HEK-APP WT and KO transfected with either EV or hGSAP. (*p-value < 0.05; ** < 0.01; ***<0.001)

Collectively, these results indicate 1) that GSAP knockout reduces γ-secretase activity towards APP but not Notch, 2) that the level of endogenous GSAP in the WT cells is saturated to modulate γ-secretase activity, and 3) that exogenous overexpression of GSAP cannot further enhance γ-secretase cleavage in WT cells.

To better understand the effect of GSAP on γ-secretase, we measured the kinetics of γ-secretase in membrane fractions prepared from four cell lines: HEK GSAP WT and HEK-GSAP KO cells transfected with EV or hGSAP. First, we found that Km and Vmax for APP of HEK-APP WT are 0.30 μM and 123.99 AU/μg/min, respectively, and for GSAP-KO are 0.25 μM and 82.034 AU/μg/min, respectively (Fig. 3A and B). Second, transfection of hGSAP in the KO cells increased Vmax (αLISA arbitrary unit) without modifying the Km value (μM) (Fig. 3B and C) but had no effect on HEK GSAP WT γ-secretase. Finally, γ-secretase from all four cell lines had similar Km and Vmax values for Notch substrate regardless of WT or KO and hGSAP transfection (Fig 3D-F). These results indicate that GSAP does not change the binding of γ-secretase to APP substrate, but rather alters the Vmax. Moreover, GSAP specifically modulates γ-secretase for Aβ production, but not Notch1 cleavage.

**Figure 3.**
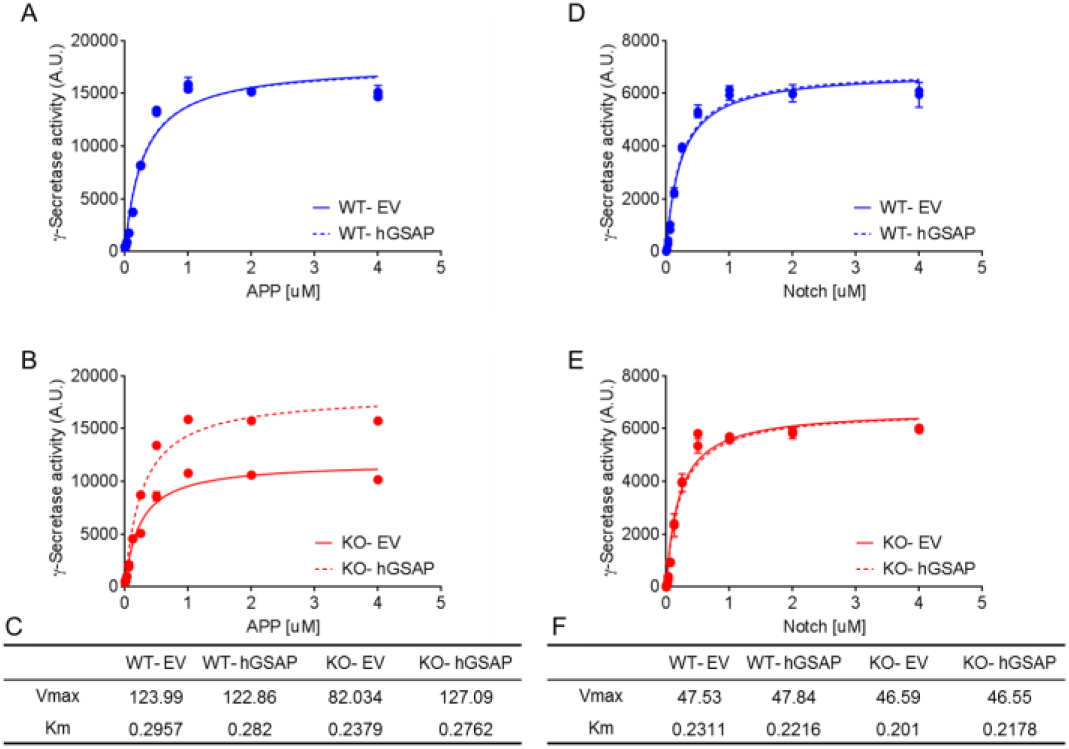
GSAP modifies γ-secretase catalytic efficiency for APP, but not for Notch. Kinetic curves fitted to Michaelis–Menten model of γ-secretase activity processing recombinant APP (A, B) or Notch (D, E) measured from membrane fraction of either HEKAPP GSAP WT (A, D) or GSAP KO (B, E) transfected with EV or hGSAP. Vmax (A.U./μg/min) and km (μM) values calculated for APP (C) or Notch (F) substrate. The data are representative of three independent experiments.

Kinetic analysis suggests that GSAP might alter the active site of γ-secretase leading to different catalytic efficiency for APP substrates. To detect the changes in the active site in the presence and absence of GSAP, we used active site-directed γ-secretase inhibitors that directly interact with PS1-NTF and -CTF ^4,18^. L631 contains two benzoylphenylalanine (BPA) groups at its P2 and P3’ positions allowing for the photolabeling of γ-secretase (Fig. 4A) ^19^. L631 has been demonstrated to label PS1-NTF, PS1-CTF and cross-linked PS1-NTF and PS1-CTF together ^19^. The photolabeling efficiency depends on direct contact between the residues and the corresponding subpockets (S2 and S3’) in the active site (Fig. 4A). Any conformational changes induced by GSAP, which alter the distance or orientation between a subpocket and the photoprobe, may lead to different cross-linking efficiencies ^18^. Four cell type membranes (WT-EV, WT-hGSAP, KO-EV and KO-hGSAP) were photolabeled with L631. Labeled species were isolated by streptavidin beads and analyzed by PS1-NTF and PS1-CTF antibodies. L631 photolabels both PS1-NTF (Fig 4B upper panels, ~34 kD band) and PS1 CTF (Fig 4B lower panels, ~20 kD band) in membranes from all four cell lines. One striking difference is that L631 doesn’t cross-link PS1-NTF and PS1-CTF together (a ~55 kDa band) in GSAP deficient cells (KO-EV) (Fig. 4B) and the labeling of the 55 kDa band can be restored by the re-expression of GSAP (KO-hGSAP). In the presence of GSAP, we propose that PS1-NTF and PS1-CTF align in a specific conformation in which L631 can cross-link the two fragments, resulting in higher γ-secretase activity for Aβ production. However, when GSAP is absent, the conformation of the active site is transformed into a different form, which does not allow for L631 to cross-link the two fragments (Fig. 4C) and results in lower γ-secretase activity for Aβ production, but not for Notch1 cleavage.

**Figure 4.**
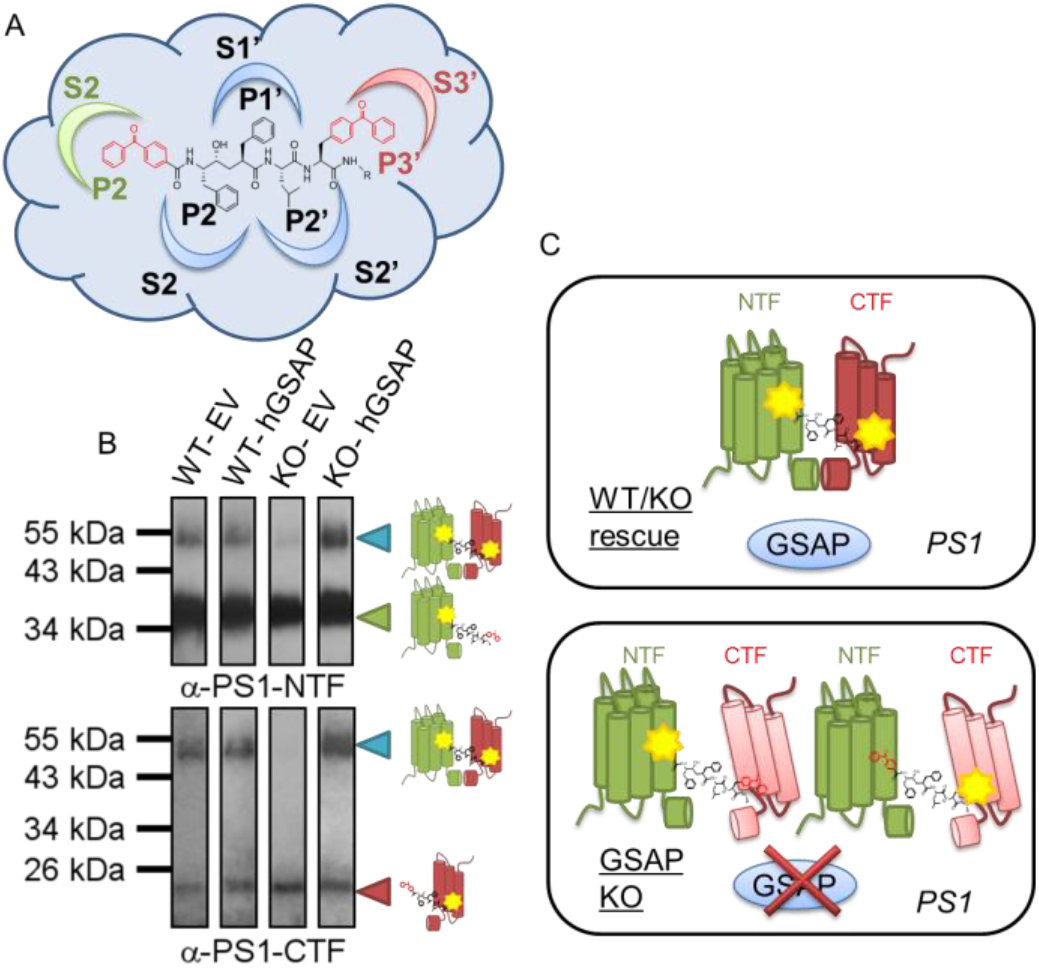
GSAP modifies the active site conformation of PS1. (A) Structure of photoprobe L631 with two BPA groups embedded in P2 and P3’ which is photoactivable and can detect conformational changes corresponding to γ-secretase active site subpocket S2 and S3’. (B) Western blot analysis with antibodies for PS1-NTF (upper panel) and PS1-CTF (lower panel) of the photolabeling efficiency by L631 in HEKAPP WT or KO transfected with EV or hGSAP. (C) Schematic representation of GSAP modification of PS1: In the presence of GSAP, PS1 NTF and PS1 CTF are aligned in a specific confirmation where the L631 probe can photolabel and cross-link both together, but when GSAP is absent this active site confirmation is modified and the L631 crosslinking is absent.

Although the effect of GSAP on the production of Aβ has been established, its role in the activation and specificity of γ-secretase has been controversial ^20^. In the present study, we show that knockout of GSAP cell lines using CRISPR/CAS9 reduces Aβ production and γ-secretase activity. We found that GSAP knockout decreases γ-secretase activity towards APP, but not Notch substrate *in vitro*. Moreover, the reduction of Aβ and γ-secretase activity was rescued by overexpressing human full-length GSAP in the knockout cells. Furthermore, our photolabeling studies indicate that γ-secretase has at least two conformations modulated through GSAP. Switching between these two forms can affect γ-secretase activity for Aβ production, but not Notch1 cleavage. When GSAP is present in the cells, like in the WT or KO transfected with GSAP (Fig. 5A), PS1 adopts a native conformation allowing for the normal processing of APP and Notch. In this native conformation, PS1-NTF and -CTF are in a specific orientation allowing for the L631 photoprobe to cross-link the two fragments. However, KO of GSAP results in a different PS1 conformation, associated with a reduction of γ-secretase activity for APP, leading to reduction in Aβ secretion (Fig. 5B). Our studies show that GSAP directly affects the conformation of the active site, providing proof that GSAP directly regulates γ-secretase activity and specificity. Our findings provide a route to the investigation of γ-secretase modulation and ultimately to the development of therapeutic agents for AD.

**Figure 5.**
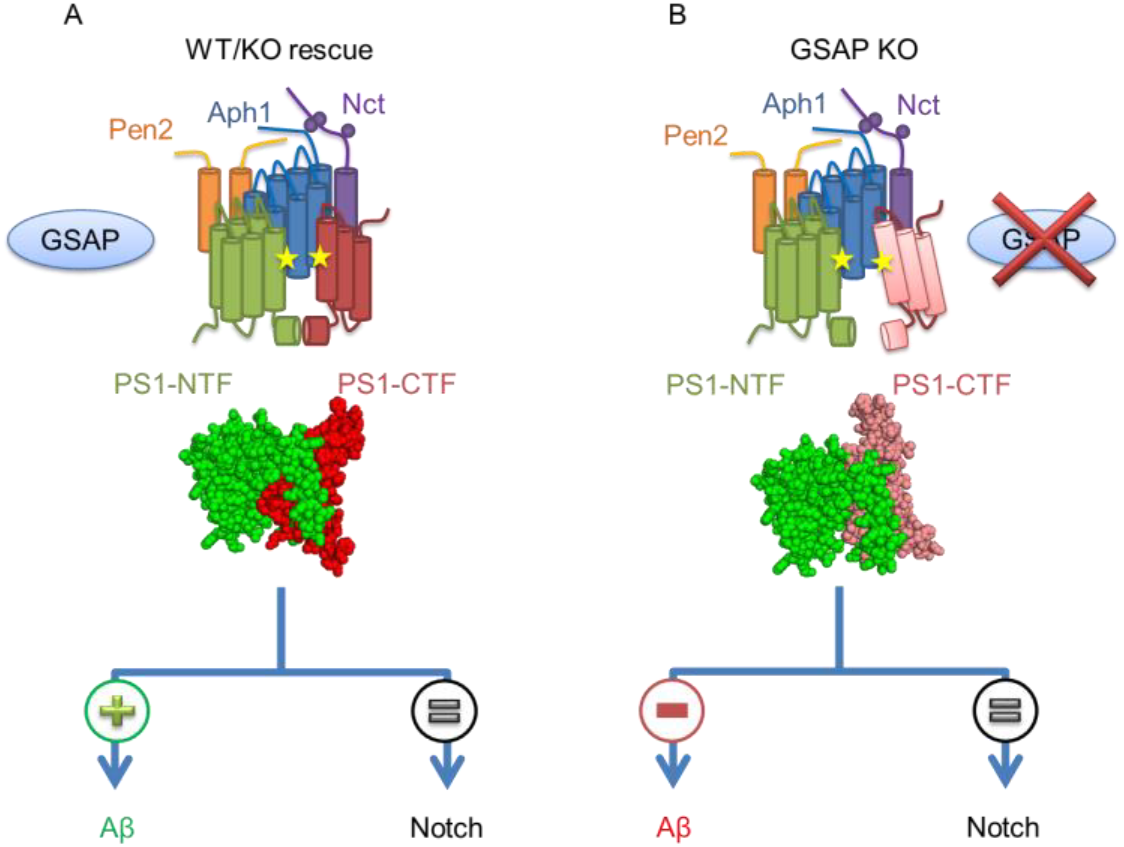
Proposed model of GSAP modulation of γ-secretase activity. γ-Secretase complex presented as transmembrane rods containing PS1-NTF (green), PS1-CTF (red), Nicastrin (purple), Aph1 (blue) and Pen2 (orange) in the present of GSAP (blue sphere) in WT or GSAP rescue with induced PS1 conformation which leads to γ-secretase activity for both APP and Notch. When GSAP is absent (B) PS1 adopts a different conformation, which leads to a decrease in APP processing and a reduction in Aβ secretion, but not in Notch processing.

## Materials and Methods

### Cell Culture

HEK-APP cell lines were cultured in DMEM supplemented with 10% FBS and 1% penicillin and streptomycin. Human neuroblastoma SH-5YSY cell lines were grown in MEM/F-12 supplemented with 10% FBS and 1% penicillin. Transfection was done using Lipofectamine® LTX with Plus™ Reagent according to manufacturer’s instructions.

### CRISPR/CAS9 GSAP KO generation and isolation

Human GSAP CRISPR/CAS9 plasmid with gRNA targeting exon 16 (CATTGCCCTTTACAGTCATT) was design and cloned into PX459 by MSKCC RNAi core facility. HEK-APP or SH-5YSY cells were transfected and and selected with 2 μg/ml Puromycin. Single clones were isolated and analyzed by DNA sequencing of GSAP exon 16. Both HEK-APP and SH-5YSY hGSAP KO clones contain a single nucleotide deletion, which creates early termination.

### RNA isolation and real-time RT-PCR

Total RNA was isolated with the Qiagen RNeasy mini kit according to manufacturer’s protocols. RNA (1 μg) was reversely transcribed to cDNA using the Superscript III 1st strand synthesis kit (Invitrogen). qRT-PCR analysis was performed with designated cDNA samples, Taqman Gene Expression Assay (Applied Biosystems). All real-time quantitative PCR was performed in triplicates on the Fast 7500 Real-time PCR system (Applied Biosystems). Taqman primers: hGSAP (Hs01383759_m1), and ribosomal 18S (Hs03003631_g1) from Applied Biosystems. Relative quantitation between samples was analyzed the ΔΔCT method.by

### Meso Scale Discovery (MSD)

Secreted human Aβ species was detected using MSD multiplex (6E10) from cell culture media 48-hour post transfection according to manufacturer’s instructions.

### Western Blot and antibodies

Cells were lysed in RIPA buffer (50mM Tris, pH8.0, 150nM NaCl, 0.1% v/v Nonidet P-40, and 0.5% wt/v deoxycholic acid) containing protease inhibitor cocktail. Protein concentration was determined by the DC assay kit (Biorad). Antibodies used for Western blot are as follows: Both PS1-NTF and Nct antibodies were from our laboratory, PS1-CTF (Millipore, MAB5232), Aph1a (Invitrogen, 38-3600), Pen2 (Abcam, 18189), APP (Millipore, MABN10) and HA (Abcam, 18181).

### γ-Secretase activity assays

The exo-cell assay was performed as described ^17^. Briefly, cells were seeded in 96-well culture plates for 24 hours, media then were removed and washed with PBS. Next, Sb4 substrate (1 μM) or NTM2 substrate (0.4 μM) was added and incubated in PIPES Buffer (50 mM PIPES, pH 7.0, 150 mM KCl, 5 mM CaCl2, 5 mM MgCl2) and 0.25% CHAPSO detergent at 37°C for 3 hours. γ-Secretase products were detected by AlphaLISA methods using G2-10 or SM320 antibodies for Aβ40 or NICD, respectively ^18^. Activity readout was expressed as arbitrary AlphaLISA units. Specific activity was normalized to protein concentration. Cell membrane preparation and g-secretase assays were described previously ^18,21,22^

### Activity-based photoaffinity labeling

Active site-based photoaffinity labeling was performed with cell lines in 12-well tissue culture dish with 10 nM of L631 in Phosphate buffer saline (PBS) pH 7.4, 0.25% CHAPSO. Photolabeling experiments were carried out as described previously ^4 19^. Reaction mixtures were solubilized with RIPA buffer for 1 hour at room temperature and streptavidin beads were added to capture labeled PS1 at 4°C overnight. Beads were washed and eluted by boiling in 2X Laemmli sample buffer. Ensuing samples were resolved on SDS-PAGE followed by western blotting with PS1-NTF or PS1-CTF antibody.

## End Matter

### Author Contributions and Notes

E.W, Y.M.L and P.G designed research, E.W, G.P.L, J.C and P.X performed research, E.W and G.P.L analyzed data; and E.W, Y.M.L and P.G. wrote the paper.

The authors declare no conflict of interest.

This article contains supporting information online. <if true>

## Acknowledgments

This work was supported by NIH grant R01NS096275 (YML), RF1AG057593 (YML), R01AG047781 (PG), the Fisher Center for Alzheimer’s Research (PG), and the JPB Foundation (YML, PG). GPL is supported by the training grant 5T32GM073546. Authors also acknowledge the MSK Cancer Center Support Grant/Core Grant (Grant P30 CA008748), Mr. William H. Goodwin and Mrs. Alice Goodwin and the Commonwealth Foundation for Cancer Research, the Experimental Therapeutics Center of MSKCC, and the William Randolph Hearst Fund in Experimental Therapeutics.

